# The P50 predicts conscious perception under tactile but not electrical somatosensory stimulation in human EEG

**DOI:** 10.1101/2025.02.28.640811

**Authors:** Jona Förster, Giovanni Vardiero, Till Nierhaus, Felix Blankenburg

**Affiliations:** Neurocomputation and Neuroimaging Unit, Freie Universität Berlin, 14195 Berlin, Berlin, Germany; Berlin School of Mind and Brain, Humboldt-Universität zu Berlin,10117 Berlin, Berlin, Germany; Charité – Universitätsmedizin Berlin, 10117 Berlin, Berlin, Germany

**Author notes:** Corresponding author: Jona Förster, FU Berlin, Habelschwerdter Allee 45, 14195 Berlin, Berlin, Germany.

**Keywords:** electrical stimulation, mechanical stimulation, somatosensory awareness, EEG, post-perceptual processes, event-related potentials

## Abstract

Past research has not reached consensus about the neural correlates of conscious somatosensory perception. Different studies have identified ERP components at various latencies as predictors of somatosensory detection, but it is still largely unclear which factors are responsible for this variation. Here, for the first time we directly compare the event-related potential correlates of stimulus detection under tactile versus electrical peri-threshold stimulation in a between-groups design, while controlling for task-relevance and post-perceptual processes with a visual-somatosensory matching task. We show that the P50 component predicts conscious perception under tactile, but not electrical stimulation: while electrical stimulation evokes a P50 already for subliminal stimuli, there is no subliminal P50 for tactile stimulation. In contrast, the P100 and N140 components robustly predict conscious detection in both stimulation groups. Tactile stimulation produced a clearly separable N80 component for detected stimuli, that appeared as part of the N140 under electrical stimulation. The P300 predicted conscious detection even though we controlled for post-perceptual processes, suggesting that it partly reflects aspects of subjective experience. Our results demonstrate that the neural correlates of somatosensory detection depend on the type of stimulation used, limiting the scope of previous results achieved using electrical stimulation.

## Introduction

Long before the seminal paper by Crick and Koch (1990) initiated the modern field of neural correlates of consciousness (NCCs) research, electrophysiological studies of conscious perception in the somatosensory domain had already debated whether early evoked potentials might constitute candidates for somatosensory NCCs. Benjamin Libet’s finding that peripheral stimulation at intensities below the threshold of detection elicited subdurally recordable responses in primary somatosensory cortex (SI) suggested that the earliest SI activity is not sufficient for conscious perception (Libet, 1993; Libet et al., 1967). This result has later been replicated (Ray et al., 1999), and more recent studies have confirmed the existence of subthreshold potentials recordable even from the scalp (Forschack et al., 2017, 2020; Nierhaus et al., 2015). Both sub- and supraliminally, the amplitude of early event-related potential (ERP) components has often been found to covary with physical stimulus parameters. In particular, the N20, believed to be the first cortical component of the somatosensory-evoked potential (SEP) reflecting the first volley of incoming peripheral activity (Desmedt & Tomberg, 1989; Tiihonen et al., 1989), has been shown to increase linearly with stimulus intensity (Forss & Jousmäki, 1998; Jousmäki & Forss, 1998; Mima et al., 1998), and the same has been found for the P50 (Forschack et al., 2020; Nierhaus et al., 2015; Schröder et al., 2021). Both the N20 and the P50 and their magnetoencephalographic (MEG) analogues have consistently been shown to originate in SI, with area 3b as main contributor to the N20 (Allison et al., 1992; Forss, Hari, et al., 1994; Mauguière et al., 1997), and areas 1 and 3b as main contributors to the P50 (Allison et al., 1992; Jones et al., 2007).

In contrast, conscious experience is better tracked by later potentials in NCC studies using somatosensory threshold detection (Auksztulewicz et al., 2012; Forschack et al., 2020; Nierhaus et al., 2015; Schröder et al., 2021; Zhang & Ding, 2009) or masking paradigms (Schubert et al., 2006). An effect of conscious perception on the N80 component has sometimes (Auksztulewicz et al., 2012; Auksztulewicz & Blankenburg, 2013), but not always (Schröder et al., 2021; Schubert et al., 2006) been found in studies applying electrical stimulation. Using mechanical stimulation in threshold detection tasks, Jones et al. (2007) found an effect of detection in the M70 event-related field (ERF) component, while Soininen and Järvilehto (1983) reported no corresponding effect. The N80 (sometimes labeled N60, N70, or N90 due to its variable onset and peak latencies) has been described early on in SEP studies using supra-threshold stimulation, and shown to be modulated by selective attention for electrical (Desmedt et al., 1983; W. R. Goff et al., 1962; Michie et al., 1987), but more consistently for mechanical stimulation (Eimer & Forster, 2003; Guidotti et al., 2023; H. Hämäläinen et al., 1990; Schubert et al., 2008; Taylor-Clarke et al., 2002; Zopf et al., 2004). Furthermore, both with electrical (Ai & Ro, 2013; Schubert et al., 2006) and mechanical (Soininen & Järvilehto, 1983) stimulation, the P100, which involves SI (Allison et al., 1992) but most likely further sources such as bilateral secondary somatosensory cortex (SII) and posterior parietal cortex as well (Forss, Hari, et al., 1994; Forss, Salmelin, et al., 1994; Mauguière et al., 1997), is sometimes predictive of perception, but this is not always the case (Schröder et al., 2021). Most consistently reported, the N140, with likely origins in bilateral SII (Auksztulewicz et al., 2012; Auksztulewicz & Blankenburg, 2013; Frot & Mauguière, 2003) and possibly frontal contributions (Allison et al., 1992) is well established as a somatosensory NCC in electrical studies (Ai & Ro, 2013; Auksztulewicz et al., 2012; Forschack et al., 2020; Schröder et al., 2021; Schubert et al., 2006; Zhang & Ding, 2009); the tactile EEG study by Soininen and Järvilehto (1983) found a comparable N190 component, and the tactile MEG study by Jones et al. (2007) identified an M135 response. Finally, the P300 (P400 in Soininen and Järvilehto, 1983) amplitude is higher for detected than for undetected stimuli, but in a recent study this effect vanished when post-perceptual processes were controlled for (Schröder et al., 2021).

However, while the P50 partly reflects initial sensory processing, top-down attentional influences on the P50 (or P40) have been reported as well both in the sub- (Forschack et al., 2017) and in the supra-threshold case (Desmedt et al., 1983; Desmedt & Tomberg, 1989; Josiassen et al., 1990), and invasive studies in awake rhesus monkeys found that the amplitude of an N1 potential around 50 ms post-stimulus varies with stimulus detection (Cauller & Kulics, 1991; Kulics, 1982). This component was proposed to correspond to the scalp-recorded N60 component in humans, likewise thought to originate from area 1 of SI (Allison et al., 1992). Moreover, human microneurographic studies suggested that a single impulse to only one or a handful of peripheral mechanoreceptors may be sufficient to induce subjective detection of that impulse (Johansson & Vallbo, 1979), a result that aligns well with the finding that weak stimulation of a single pyramidal neuron in barrel cortex suffices to elicit detection report behavior in rats (Houweling & Brecht, 2008). In humans, Soininen and Järvilehto (1983) found that the detection of mechanical stimuli delivered to the back of the left hand at subjective threshold intensity evoked a P50 component that was absent for undetected stimuli of the same intensity. Some more recent studies have likewise found early ERP or ERF differences for detected vs. undetected stimuli, at 70 ms (Hirvonen & Palva, 2016; Jones et al., 2007) or even 30 ms post-stimulus (Palva et al., 2005).

In sum, potentials in the P50 time range were found not to be predictive of perceptual experience by most, but not by all studies of conscious somatosensory detection. This raises the question what is responsible for these divergent findings. While the above-cited studies also vary in the stimulation intensities and sites, as well as different tasks used, one obvious contender for the difference-maker is the type of stimulation: whereas most studies employed electrical stimulation (usually of the median nerve at the wrist of or one or more fingers), only a small minority used mechanical tactile stimulation. And while among the studies that report early ERP/ERF differences between detected and undetected stimuli some used electrical (Hirvonen & Palva, 2016; Palva et al., 2005) and some tactile stimulation (Jones et al., 2007; Soininen & Järvilehto, 1983), the studies that reported no connection between early potential amplitudes and conscious detection invariably used electrical stimulation (Auksztulewicz et al., 2012; Forschack et al., 2020; Libet et al., 1967; Ray et al., 1999; Schröder et al., 2021; Schubert et al., 2006; Uemura et al., 2021; Zhang & Ding, 2009). Thus, there is the intriguing possibility that the NCCs of somatosensory detection might differ for tactile compared to electrical stimuli. While there are several somatosensory-evoked potential (SEP) studies comparing different stimulation types for strong supra-threshold stimuli (e.g., mechanical fingernail stimulation versus electrical stimulation: Pratt et al., 1979; pin pricks and taps versus electrical stimulation: Yamauchi et al., 1981; air-puffs versus electrical stimulation: Forss et al., 1994; Rossini et al., 1996), to our knowledge no study so far systematically compared the difference in early ERPs between detected and undetected peri-threshold stimuli for different stimulus types.

Therefore, in this study we use EEG to compare the evoked responses to mechanical versus electrical stimulation of the left index finger at various sub- and suprathreshold intensities in a between-groups design, with considerably larger sample sizes than the previous mechanical stimulation studies (Jones et al., 2007; Soininen & Järvilehto, 1983). Simultaneously, we control for possible effects of attention and post-perceptual processes such as perceptual decision-making by means of a visual-somatosensory matching task. This is important because without such control, later occurring ERP components can be mistaken for NCCs when they really reflect differences in task-relevance or response preparation (Förster et al., 2020; Pitts et al., 2014; Schröder et al., 2021). Our aim in this study was to investigate, for each of the components introduced above (N20, P50, N80, P100, N140, and P300), how its amplitude is affected 1) by the perception or lack thereof of peri-threshold stimuli, and 2) how this effect is modulated by the type of stimulation used (electrical or tactile).

## Methods and Materials

### Participants

Participants were recruited among the students of Freie Universität Berlin. All participants gave their written informed consent and declared to have no physical or psychological illness, and to be right-handed according to the Edinburgh Handedness Inventory (Oldfield, 1971). Participants first had to perform a behavioral training session in order to ensure that they had stable psychometric functions and sufficient task performance to be included in the main experiment, which took place on a separate day. Twenty-nine participants completed the mechanical stimulation experiment (tactile group). Of these, four participants were excluded from the analysis due to poor psychometric functions in the main experiment, resulting in twenty-five participants (14 female, 11 male, age range 19-32 years) that were included in the analyses. The electrical stimulation experiment was completed by twenty-eight participants (electrical group), of which three were excluded for the same reason as in the tactile group, again resulting in twenty-five participants (16 female, 9 male, age range 21-35 years). Participants received compensation in the form of either money or course credits. The study was approved by the local ethics committee of the Freie Universität Berlin (003/2021) and conducted in accordance with the Declaration of Helsinki.

### Experimental Design

Participants performed a two-alternative forced-choice detection task via a visual-somatosensory matching task (Fig. 1) that has already been employed in several previous studies (Schröder et al., 2019, 2021). Participants were seated in front of a computer screen and their eye movements were recorded with an eye tracker (SMI RED-m remote, 120 Hz, Sensomotoric Instruments, Teltow, Germany). Every trial started with the appearance of a medium brightness gray fixation disk at the center of a black background. Participants were then presented with either mechanical (tactile group) or electrical (electrical group) stimuli, delivered at 1 of 10 different intensities to their left fingertip. The intensities were chosen to sample the entire individual psychometric function, which served to maintain participants’ attention and interest in the task and prevent them from mere guessing, and enabled us to identify slight shifts of the detection threshold over the course of the experiment and to monitor task performance. Simultaneous to the onset of stimulation, a visual matching cue was presented for 800 ms. The matching cue consisted of a change in brightness of the gray fixation disk to either white or dark gray, signifying target presence or absence, respectively. After each stimulus presentation, the fixation disk returned to medium brightness for 300 ms, and two colored disks appeared for 900 ms on the left and right sides on the screen (counterbalanced across trials). The colors coded for “match” and “mismatch” (counterbalanced across participants), and participants responded by directing their gaze to the disk that corresponded to their experience (match or mismatch of somatosensory experience and visual cue). This match-mismatch procedure served to decorrelate somatosensory target detection from overt reports, while using saccades instead of button presses to respond ensured that stimulus-evoked electrophysiological activity from somatosensory regions could not be influenced by response-related activity from the hand region of the adjacent motor cortex. When participants gave their response in time, the chosen response cue briefly increased in size; when they were too slow (>0.9 s), the gray fixation disk in the center of the screen briefly turned red, signaling a missed trial. Individual trials were separated by intertrial intervals (ITIs) between 0.7 and 1.3 s.

**Figure 1.**
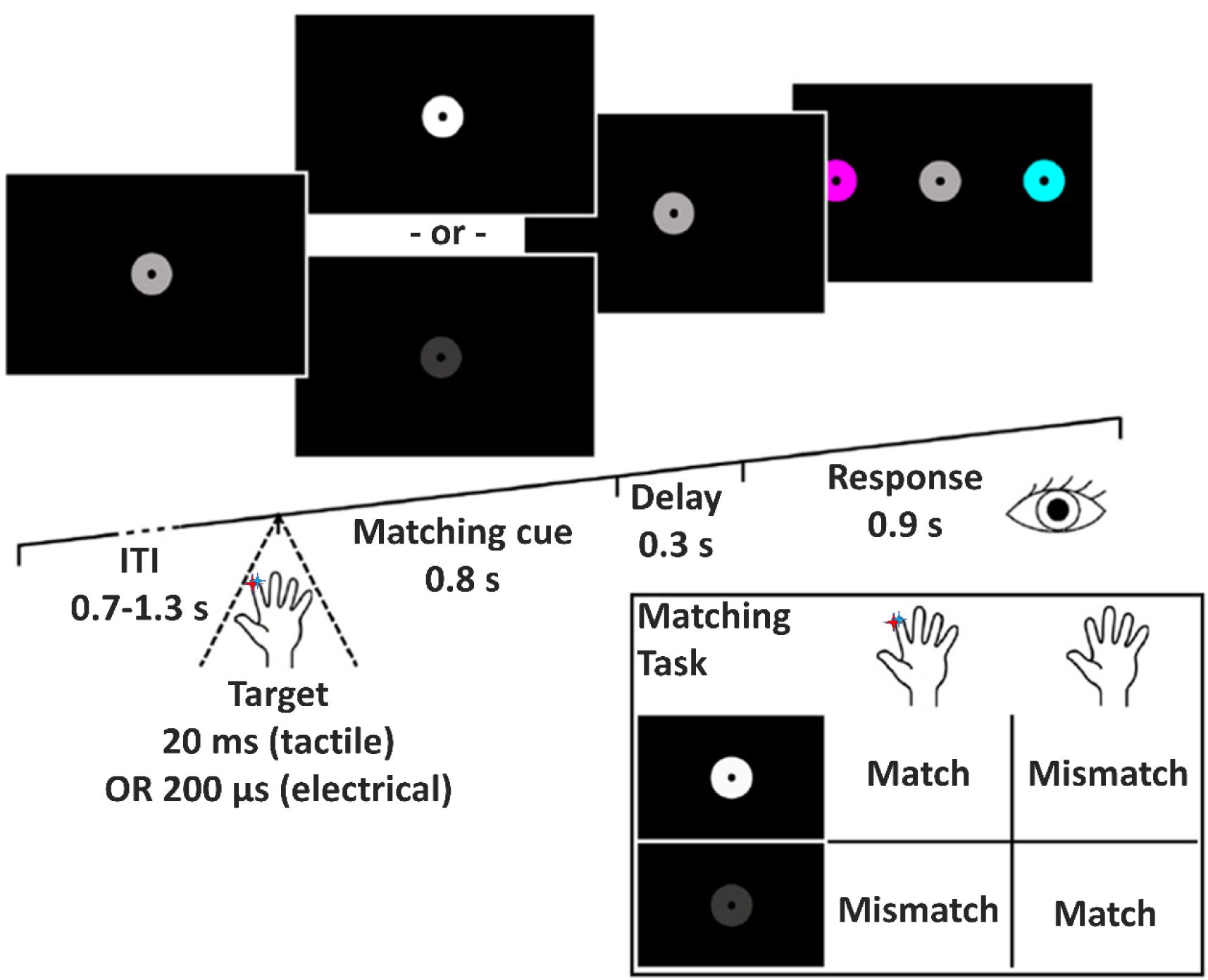
Experimental Design. After a jittered ITI, participants received a mechanical impulse (tactile group) or an electrical pulse (electrical group) to the pulp of the distal phalange of their left index finger at 1 of 10 individually calibrated intensities on each trial. Simultaneously, the gray fixation disk turned into a visual matching cue by changing its brightness, signaling either target presence (white) or target absence (dark gray). Participants then compared their percept (detected vs. not detected) to the visual cue and decided whether their somatosensory experience matched the meaning of the cue (according to the box on the lower right side of the Figure). Following a brief delay, participants reported their decision by saccading to one of two color-coded response cues presented at the sides of the screen.

#### Stimuli

The mechanical stimuli were impulses with 20 ms duration (the shortest time that would yield stable psychometric functions when piloting the study) that were delivered to the pulp of the left index finger’s distal phalange using a single pin of a piezoelectric Braille display controlled by a programmable stimulator (Piezostimulator, QuaeroSys, St. Johann, Germany). The Braille module was taped to the finger firmly enough to prevent the pin position from changing, but gently enough not to cause blood flow sensations that might be confused with the mechanical stimulation. Participants in the tactile group wore earplugs to prevent auditory perception of the pin motion. Stimulus intensity was controlled via the pin’s protrusion (0-1.5 mm, controlled in arbitrary units [a.u.] of 0-4096). The electrical stimuli were direct current square wave pulses of 0.2 ms duration (a time commonly used in comparable studies, cf. Auksztulewicz et al., 2012; Auksztulewicz & Blankenburg, 2013; Schröder et al., 2021), delivered via adhesive electrodes (GVB-geliMED, Bad Segeberg, Germany) at the same location using a DS5 constant current generator (Digitimer Limited, Welwyn Garden City, Hertfordshire, UK). In both groups, we began by determining participants’ individual detection threshold by a brief staircase procedure: starting from an initial intensity value (tactile: 1000 a.u., electrical: 0.5 mA), this value was increased by 100 a.u. (tactile) or 0.1 mA (electrical) until the participant reported to feel the pulse. The stepsize was then halved, and the intensity decreased by the new stepsize until the participant reported not feeling the pulse anymore. The stepsize was then again halved, and the intensity increased by the new stepsize, and so on for three up- and three down-progressions in total. Starting from these values, participants’ psychometric functions were estimated in order to accommodate between-subject variation in detection thresholds and criteria. Participants received 15 intensities (20 repetitions per intensity, leading to 300 trials in total), linearly spaced around their initial detection threshold. After each stimulus presentation, they were required to respond via keyboard whether they had detected the pulse. A logistic function with two parameters (detection threshold and slope at threshold) was then fitted to the data (estimated 1%, 50%, and 99% detection thresholds: T01 = 534 ± 159 a.u., T50 = 834 ± 118 a.u., T99 = 1134 ± 223 a.u. in the tactile group, and T01 = 1.09 ± 0.4 mA, T50 = 1.46 ± 0.42 mA, T99 = 1.82 ± 0.52 mA in the electrical group; all descriptive statistics are reported as mean ± SD, except when otherwise noted). Based on these parameters, 10 different equally spaced intensity levels were determined and used in the main experiment. Based on previous studies (Schröder et al., 2019, 2021), the trial numbers within each intensity level followed a normal distribution, such that most trials occurred with an intensity close to the individual detection threshold (intensity levels 5 and 6: 32 trials/run each), and relatively few trials with intensities far from threshold (intensity levels 1 and 10: 8 trials/run each). Stimuli were presented in MATLAB 2013a (The MathWorks) via the Psychophysics toolbox (Brainard, 1997).

### Experimental procedure and EEG recording

All participants performed a behavioral training session prior to the experiment and on a separate day to ensure that they had stable psychometric functions, and were invited to the EEG recording only if they reached at least 90% accuracy in a training run with only sub- and suprathreshold stimulation, demonstrating comprehension of the task and ability to perform it correctly at low error rate. EEG data were recorded from 64 active electrodes positioned according to the extended 10-20 system (ActiveTwo, BioSemi, Amsterdam, Netherlands) with 2048 Hz sampling frequency. Vertical (vEOG) and horizontal (hEOG) eye movements were recorded with four additional electrodes. In the tactile group, eleven participants completed seven runs of 200 trials each (∼10 min per block), resulting in a total number of 1400 trials per participant. The remaining fourteen participants in the tactile group completed only six runs (1200 trials), due to fatigue in one or more runs. In the electrical group, twenty participants completed seven blocks, and five participants completed only six runs. After the main experiment, a localizer run with suprathreshold stimulation at 2 Hz frequency (jittered with ±10 ms, uniform distribution) was recorded (tactile group: 800 trials, electrical group: 600 trials) to enable delay correction (see below). The intensity was set to be several times above detection threshold, but below motor threshold (tactile group: 4000 a.u., electrical group: 4.05 mA ± 0.77 mA).

### Data analysis

#### Behavior

To visualize the distribution of detection thresholds across participants, we fitted logistic functions to the behavioral data of each run and averaged the estimated slope and normalized threshold parameters for each participant, resulting in one mean psychometric function per participant (Fig. 2). Estimated detection probabilities < 10% for intensity level 1 and > 90% for intensity level 10 were defined as inclusion criteria to minimize the possibility of incomplete sampling of individuals’ psychometric functions (due to shifts in detection thresholds, response criteria, or erroneous reports). Slope differences between the two groups and reaction time differences between hits and misses within groups were tested using a Bayesian paired-sample t-test equivalent (Krekelberg, 2022); we report Bayes factors in favor of a difference (BF10). To test whether the matching task was successful in dissociating somatosensory target detection (hits vs. misses) from overt reports (match vs. mismatch), we performed Bayesian tests of association (the Bayesian equivalent to a chi-square test; cf. Albert, 1997) for all participants and report Bayes factors in favor of the null hypothesis (BF01). Following the recommendations by Kass and Raftery (1995), we consider 1 ≤ BF < 3 negligible, 3 ≤ BF < 20 positive, 20 ≤ BF < 150 strong, and 150 ≤ BF very strong evidence.

**Figure 2.**
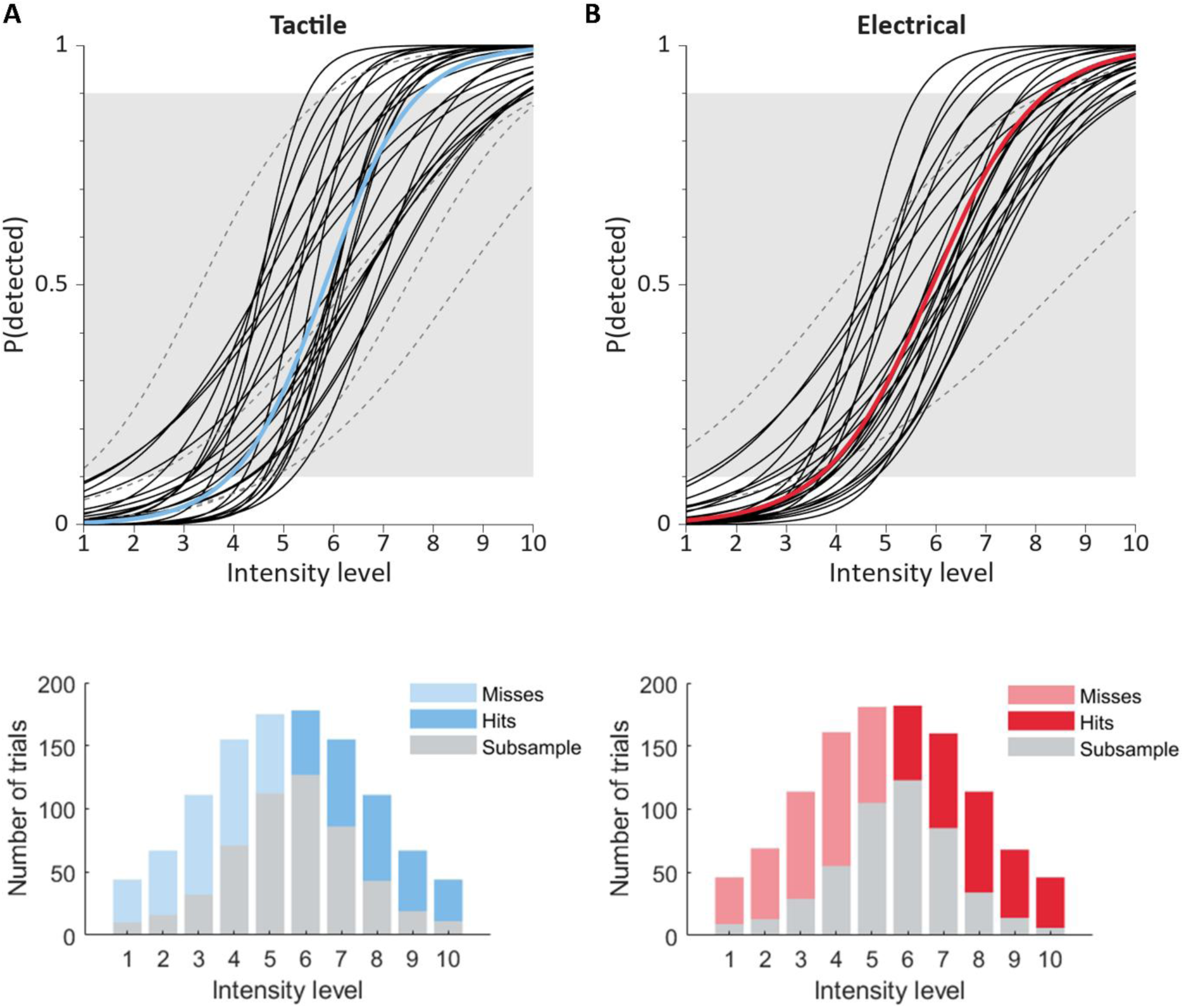
***A:** Upper panel:* Normalized mean psychometric function in the tactile group (n=25). Black lines indicate the individual psychometric functions included in the final sample, averaged over runs and normalized across participants to intensity levels 1-10. Blue thick line denotes the group average. Gray dashed lines indicate participants whose detection probabilities at minimum and maximum intensity levels fell outside the required margin of <10% and >90% (white background) and were thus excluded from the analysis. *Lower panel:* Histograms of trial numbers before and after subsampling for the tactile group. Gray color denotes the trial numbers after subsampling, which are by definition the same for hits and misses within each intensity level. ***B:*** Same as ***A***, but for the electrical group (n=25). Red thick line in the upper panel denotes the group average.

#### EEG preprocessing

Preprocessing and all analyses were performed using SPM12 for EEG (www.fil.ion.ucl.ac.uk/spm) and custom MATLAB scripts. The data were high-pass filtered at 0.01 Hz, notch-filtered between 48-52 Hz, down-sampled to 512 Hz, and re-referenced to common average. Eye blinks were removed from the data using adaptive spatial filtering based on individual blink templates computed from the vEOG (Ille et al., 2002). The data were then epoched from -50 to 600 ms relative to stimulation onset. All epochs were visually inspected for artifacts. Bad channels (containing >20% bad trials) were interpolated on a run-by-run basis when only specific runs were affected, or completely when all runs were affected (tactile group: 3.4 ± 1.9 channels, electrical group: 4 ± 2 channels interpolated in at least one run). The remaining artifactual trials were removed (tactile group: 11.4 ± 3.3%, electrical group: 13.4 ± 3.9%). The data were then low-pass filtered at 40 Hz and baseline-corrected using a baseline from -50 to -5 ms. Because mechanical stimulation with the Quaerosys stimulator is slightly delayed compared to electrical stimulation with the DS5 and we used different stimulus durations in the two groups, we used the localizer data to measure the peak latency of the earliest physiological component that could clearly be discerned in the localizer data, which was the P50 in the CP4 electrode (tactile group: 50.78 ms; electrical group: 42.97 ms, and corrected all EEG data of the tactile group by that delay (7.81 ms). Following the SPM standard procedure, the preprocessed, epoched channel data were linearly interpolated into 32×32×334 voxel (scalp space x intra-trial samples) 3-D images per trial and smoothed with a 8 by 8 mm full-width half-maximum Gaussian kernel.

### EEG data analysis

To test the influence of conscious detection (hit vs. miss) and stimulation (mechanical vs. electrical) on the various ERP components, we implemented a factorial between-groups design in SPM and investigated the main and interaction effects. We performed mass-univariate general linear model (GLM) analysis of the entire sensor-time space of a report-matched subsample of the data. The advantages of our experimental design come at the cost of introducing a degree of collinearity between stimulation intensity and conscious perception. To circumvent this problem, we defined a subsample of the data by matching the number of “hit” and “miss” trials in each of the ten intensity categories for each participant. In effect, this means discarding trials mostly at the lower (where misses dominate) and higher (where hits dominate) intensities, and keeping mostly trials of the intermediate intensities (where hits and misses are most balanced). The mean number of trials per subsample across participants was 525 ± 96 in the tactile group, and 475 ± 76 in the electrical group (Fig. 2). To ensure that our analysis is not biased by the peculiarities of a particular subsample, we randomly drew 100 such subsamples, performed all subject-level classical GLM analyses on each of them, averaged the results, and took only this average to the group level.

On the subject level, we performed two-sample t-tests between hit and miss trials. Due to the large number of models to be estimated (2 x 25 participants x 100 subsamples), these analyses were carried out on the high-performance cluster (HPC) of the Freie Universität Berlin (Bennett et al., 2020). For each subject, we computed the average over subsamples of the resulting beta estimates. On the group level, we used SPM’s flexible factorial specification to estimate a 2×2 design with the factors group (levels: tactile/electrical) and detection (levels: hit/miss). Based on the ERP plots, we investigated t-contrasts for the main effects hit > miss (for the P50, P100, and P300 components) and miss > hit (for the N20, N80 and N140 components), as well as for the two directions of interaction. For the main effect of group (tactile vs. electrical), we formulated an F-contrast to get an overall impression of the spatiotemporal group differences. We report results thresholded with p < 0.05 FWE-corrected at the peak level, using the random field theory-based approach to multiple comparisons correction as implemented in SPM.

## Results

### Behavior

Participants detected 47.77 ± 9.65% of the targets in the tactile group and 46.1 ± 9.08% in the electrical group. As expected by design, experience varied the most on trials with intermediate intensities, leading to sigmoidal psychometric curves (Fig. 2). We found no evidence for a difference in slopes between the two groups (BF10 = 0.95). We found strong evidence for a difference in reaction times between hits and misses in both the tactile (hits: 309.54 ± 40.46 ms, misses: 317.72 ± 43.81 ms, BF10 = 197.87) and the electrical group (hits: 316.72 ± 44.66 ms, misses: 322.59 ± 44.33 ms, BF10 = 5.24). Bayesian tests of association provided positive evidence that the matching task successfully dissociated target detection from overt reports (3 < BF01 < 10 for all participants in both groups).

### Event-related potentials

The electrode regions typically affected in somatosensory detection tasks exhibited most of the characteristic ERP components we expected based on the established literature. The N20 component was not discernible with certainty and, if present at all, of very small amplitude. The P50 in contralateral centroparietal electrodes (Fig. 3), the P100 in central electrodes, the N140 in contralateral frontocentral electrodes, and the P300 in central electrodes (Fig. 4) were clearly present in both the tactile and the electrical group. In addition, the tactile group exhibited a clearly separate N80 component for hits, preceding the N140 in the same sensors (Fig. 4). Remarkably, while the P50 and P100 amplitudes in electrode CP4 were only slightly smaller for misses compared to hits in the electrical group, these components were practically absent for misses in the tactile group. The N140 in electrodes C6 and FC6 was weaker for misses than hits in both groups, but much more so in the tactile group. The electrically evoked P300 in electrode CPz reached a plateau already around 200 ms, whereas the tactile-evoked P300 began later and peaked sharply at 300 ms before it settled at about the same amplitude as in the electrical group. In both groups, the P300 amplitude was stronger for hits than misses.

**Figure 3.**
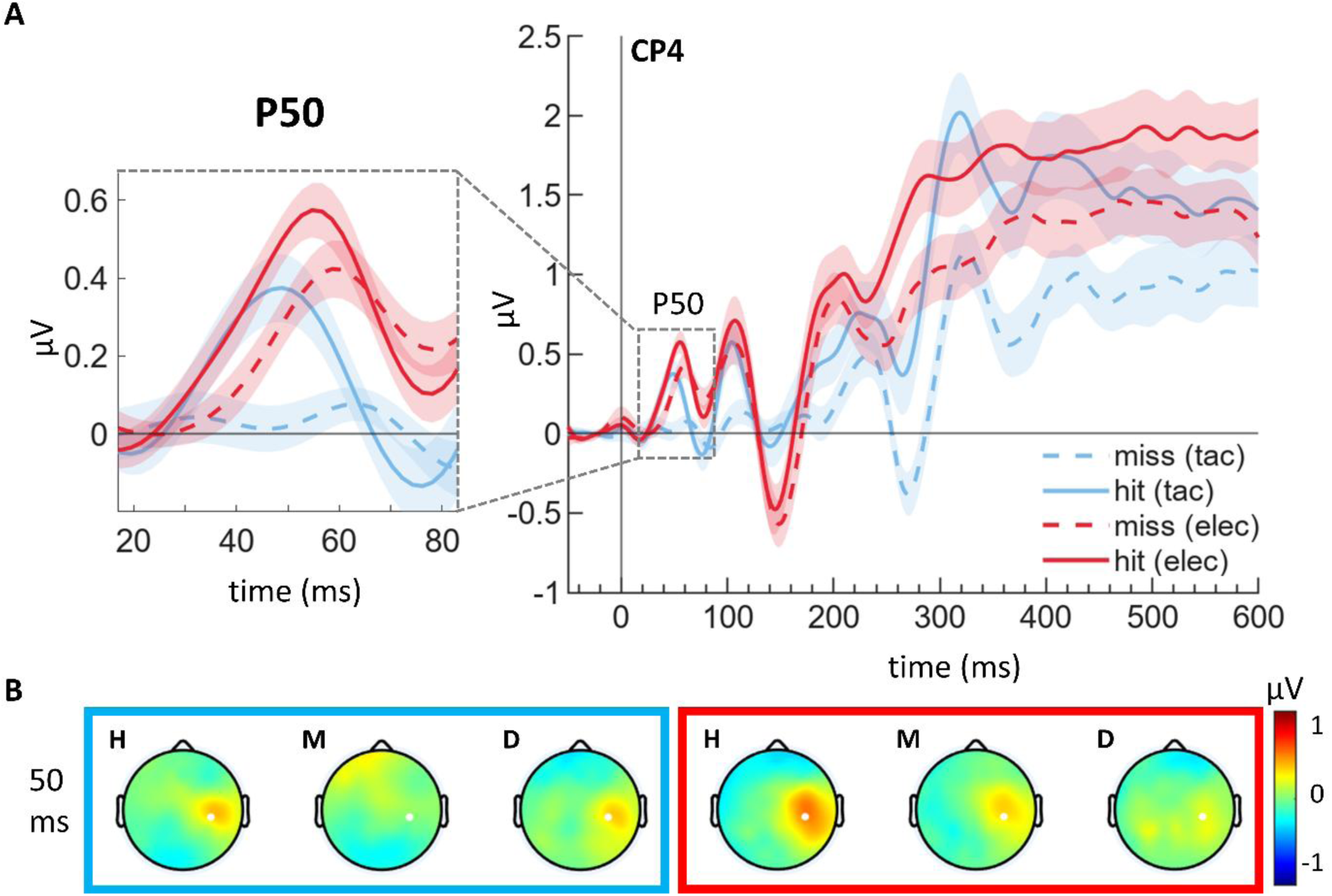
***A:*** ERP plot of electrode CP4, where the P50 component and the statistical main effect of conscious detection in this time range were most prominent. Left side shows the P50 time window magnified. ***B:*** Scalp voltage topographies at 50 ms after target presentation for the tactile (left, blue box) and electrical (right, red box) groups separately for hits (H), misses (M) and their difference (D). White dots denote the electrode location of the corresponding ERP plots, where the statistical effect was maximal.

**Figure 4.**
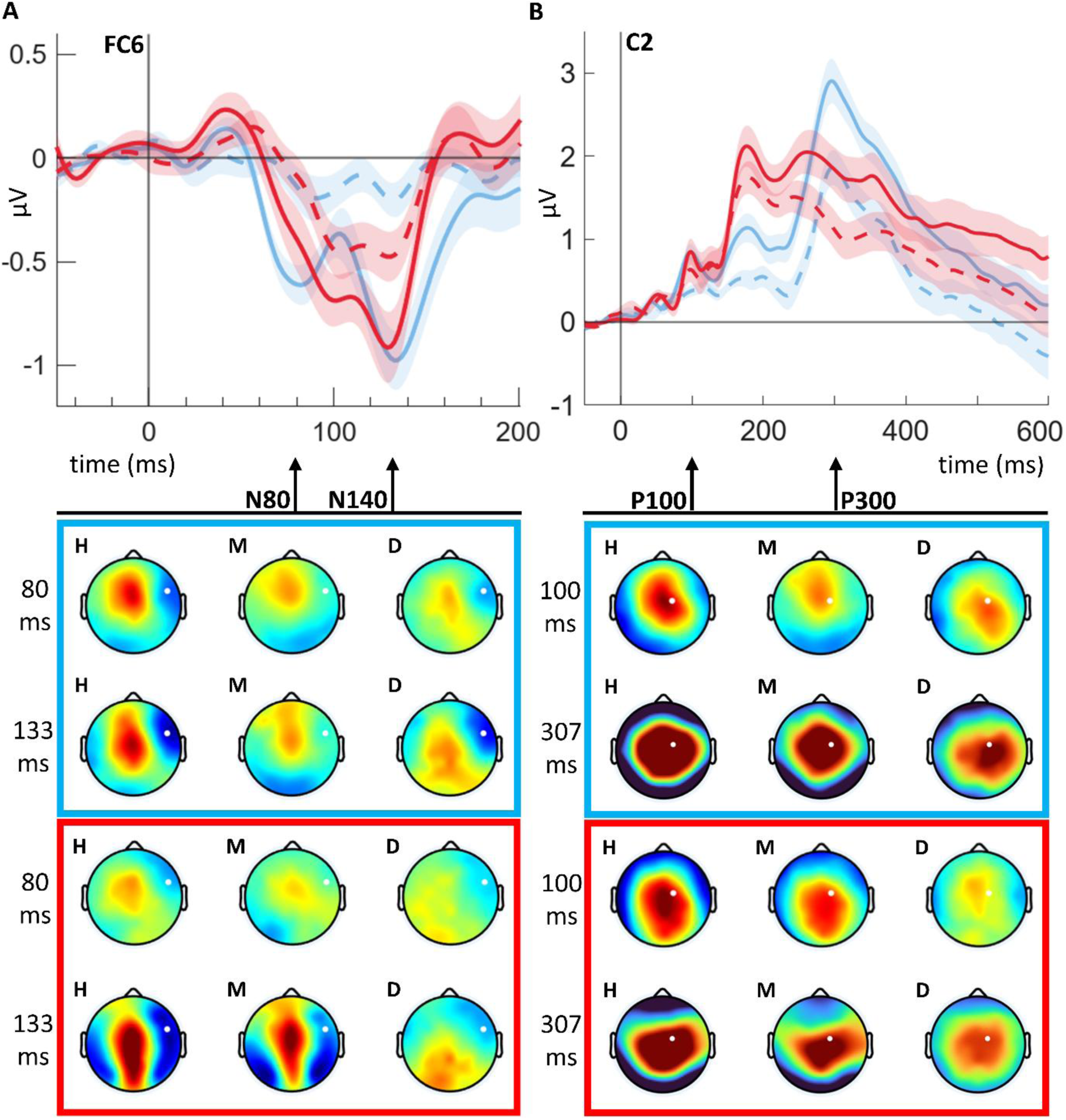
Subsample trial distributions, event-related potentials, and topographies. ***A:*** *Upper panel:* ERP of electrode FC6, where the N80 and N140 components were most prominent. Time window from -50 to 200 ms. *Lower panel:* Scalp voltage topographies at 80 ms and 133 ms for the tactile (upper box, blue) and electrical (lower box, red) groups separately for hits (H), misses (M) and their difference (D). White dots denote the electrode location of the corresponding ERP plot in the upper panel. ***B:*** *Upper panel:* ERP of electrode C2, where the P100 and P300 components were most prominent. *Lower panel:* Topographies at 100 ms and 307 ms. Color coding and white dots as in ***A***.

In the statistical analysis, the main effect of condition for the contrast hit > miss revealed three significant clusters peaking at 49 ms near electrode CP4 (p*_FWE_* = 0.004), 100 ms near electrode C2 (p*_FWE_* < 0.001), and 307 ms near C4 (p*_FWE_* < 0.001), corresponding to the amplitude differences between hits and misses in the P50, P100 and P300 components of the ERP. There were two further clusters with small sizes of 6 voxels (148 ms, p*_FWE_* = 0.034) and 2 voxels (123 ms, p*_FWE_* = 0.047) respectively. The contrast miss > hit revealed a contralateral cluster peaking at 129 ms near FC6 (p*_FWE_*< 0.001), as well as earlier at 70 ms (p*_FWE_* = 0.001) and, near FT8, at 84 ms (p*_FWE_* = 0.001), corresponding to amplitude differences in the N140 and N80 components, respectively. Further clusters included ipsilateral clusters peaking at 100 ms (p*_FWE_* = 0.009) near T7, and at 166 ms (p*_FWE_* < 0.001) near C5, as well as two later clusters peaking at 309 ms (p*_FWE_* < 0.001) near P9, and at 586 ms (p*_FWE_* < 0.001) near F7, as well as a number of smaller clusters with sizes of 48 voxels (309 ms, p*_FWE_* = 0.016), 28 voxels (221 ms, p*_FWE_* = 0.016), 25 voxels (309 ms, p*_FWE_* = 0.002), 23 voxels (229 ms, p*_FWE_* = 0.024), 22 voxels (219 ms, p*_FWE_* = 0.029), 7 voxels (102 ms, p*_FWE_* = 0.032), 6 voxels (230 ms, p*_FWE_* = 0.045), and 2 voxels (102 ms, p*_FWE_* = 0.047), respectively.

There was no effect in the N20 time range, and the amplitude of this component was very small in both stimulation modalities. The main effect of group showed spatiotemporally widespread clusters throughout the entire peri-stimulus interval from ca. 100 ms onward, indicating overall amplitude differences between the signals from the tactile and the electrical group across all sensors for most of the time. There were no interaction effects. All clusters are summarized in Table 1. To investigate which group was the main driver of the P50 main effect, we inspected the contrast hit > miss for the two groups separately. As we expected based on the ERP plot, the effect at 49 ms was more strongly driven by the tactile than by the electrical group: while the effect prevailed in the tactile group (p*_FWE_* = 0.044), there was no corresponding cluster when considering the electrical group in isolation.

**Table 1.** **R**esults (main analysis)

| **Table 1. Results (main analysis)** |  |  |  |
| --- | --- | --- | --- |
| Main effect of condition: hit > miss (t-contrast, $k > 50$ voxels) | | | |
| Peak time (ms) | Nearest electrode | Cluster size $k$ | p-value (FWE-corrected) |
| 49 | CP4 | 66 | 0.004 |
| 100 | C2 | 715 | <0.001 |
| 307 | C4 | 28210 | <0.001 |
| Main effect of condition: miss > hit (t-contrast) |  |  |  |
| 100 | T7 | 69 | 0.009 |
| 129 | FC6 | 2354 | <0.001 |
| 166 | C5 | 1391 | <0.001 |
| 309 | P7 | 489 | <0.001 |
| 586 | F7 | 21133 | <0.001 |
| Main effect of group (F-contrast): |  |  |  |
| 152 | P8 | 18507 | <0.001 |
| 166 | FCz | 45416 | <0.001 |
| 193 | Fp1 | 2024 | <0.001 |
| 295 | FCz | 12614 | <0.001 |
| 320 | F7 | 933 | <0.001 |
| 420 | Oz | 52 | 0.02 |
| 496 | T8 | 121 | 0.004 |
| 500 | Fz | 3831 | <0.001 |

Following the main analysis, we performed a series of control analyses. First, we contrasted the hits from only the intermediate intensity levels 4 to 7 against all trials from the lowest intensity levels 1 to 3. The purpose of this analysis was to provide an alternative partitioning of the experimental trials, based on the assumption that trials reported as hits for such low intensities are likely to be accidental misreports, and can therefore effectively be treated as misses. This partitioning of course contains an intensity confound, but it can nonetheless serve as an informative sanity check, in that failure to replicate the results from the main analysis would seriously call them into question. All three positive effects were replicated under these conditions: there were three clusters peaking at 51 ms (p*_FWE_* = 0.006) near CP4, at 96 ms (p*_FWE_* < 0.001) near Cz, and at 306 ms (p*_FWE_*< 0.001) near C2. The converse control analysis of the miss trials from intensity levels 4 to 7 against all trials from the highest intensity levels 8 to 10 (which are likely to have mostly been perceived) again replicated the cluster peaking at 133 ms (p*_FWE_* < 0.001) near FC6. The two other peaks of that cluster were minimally spatiotemporally shifted compared to the main analysis, to 68 ms (p*_FWE_* = 0.019) near F6, and 82 ms (p*_FWE_* < 0.001) near T8. The other clusters of this contrast were extremely similar to those of the respective main analysis, both qualitatively and quantitatively.

Next, we performed analyses to test whether there would be effects of the report type (match vs. mismatch), the direction of gaze response (left vs. right saccade), or the visual matching cue (white vs. dark). There were no effects of report type and gaze direction, but there was a significant main effect of the cue, comprising several clusters beginning at 84 ms (p*_FWE_* < 0.001) in occipito-parietal and lasting up to 420 ms (p*_FWE_* < 0.001) in fronto-central electrodes. To exclude any possible influence of this cue effect on the results fromour main analysis, we drew 100 new subsamples, this time balancing the number of white and dark cues in addition to balancing the number of hits and misses within each intensity level. This analysis reproduced all our main effects, i.e., the effects of P50 at 47 ms (p*_FWE_* = 0.009), N80 at 70 ms (p*_FWE_* = 0.002) and 84 ms (p*_FWE_* = 0.001), P100 at 102 ms (p*_FWE_*< 0.001), N140 at 123 ms (p*_FWE_* < 0.001), and P300 at 305 ms (p*_FWE_* < 0.001). This confirms that the results from our main analysis were not confounded by the visual matching cue, and increases confidence that they were genuinely driven by the difference in conscious perception.

## Discussion

In this study, we investigated how mechanical and electrical stimulation, under otherwise largely identical conditions, differentially affect the event-related potentials associated with somatosensory target detection, while controlling for the influence of attention and post-perceptual processes. We found significant main effects of detection for the P50, N80, P100, N140, and P300 components. In the case of the P50, this main effect was driven by the tactile group, as indicated by the group-specific contrasts.

*N20.* We found only a negligible N20 component in either of the two groups, and no effect of perception in that time range. In electrical finger as compared to median nerve stimulation at the wrist, the N20 component is usually greatly reduced in amplitude or even completely absent (W. R. Goff et al., 1962; Rossini et al., 1996), and in mechanical finger stimulation, it is usually not present at all (Eimer & Forster, 2003; Onishi et al., 2010; Yamashiro et al., 2019; Zopf et al., 2004). Using electrical finger stimulation, Palva et al. (2005) have found an effect of conscious detection in an ERF component already 30 ms after stimulus onset.

While this is exceptionally early (we discuss a possible explanation for this finding below), it is more likely to correspond to the P27m (for “magnetic”) reported by Tiihonen et al. (1989), or the frequently reported P35m (Forss, Hari, et al., 1994; Forss, Salmelin, et al., 1994; Jousmäki & Forss, 1998; Wikström et al., 1996), which were recorded using supra-threshold stimulation. These positive components are clearly distinct from the N20(m) in terms of both source localization and behavior in response to experimental manipulations such as stimulus intensity and ISI (Jousmäki & Forss, 1998; Wikström et al., 1996), while the P35m and P60m share very similar characteristics (Wikström et al., 1996). Similarly, in an SEP study, Desmedt and Tomberg (1989) have identified a “cognitive P30” that, unlike the N20, was enhanced for attended compared to unattended stimuli. In all likelihood then, the SEF effect at 30 ms reported by Palva et al. (2005) is rather related to the P30 or even the P50, but not to the N20 SEP component.

*P50.* The P50 component in the electrical group was present for both hits and misses. Although it was slightly weaker for misses than hits and contributed to the main effect of detection at 49 ms, this effect was not present when considering the electrical group in isolation. In striking contrast, while there was a P50 for detected targets in the tactile group, that component was completely absent when targets were not detected in this group, and this effect prevailed even for the tactile group alone. Regarding the P50, this result is in line with the finding by Soininen and Järvilehto (1983), who had mechanically stimulated the hairy skin of the back of the hand and found P50, P100, N190, and P400 only for detected trials. It is also in line with almost all threshold detection studies using electrical stimulation (both of fingers and of the MN), which found no differences in the P50 time range for hits compared to misses (Auksztulewicz et al., 2012; Schubert et al., 2006; Uemura et al., 2021; Zhang & Ding, 2009), with some of them showing that P50 amplitudes vary with physical stimulus intensity instead (Forschack et al., 2020; Nierhaus et al., 2015; Schröder et al., 2021). The same linear scaling effect has been reported for early magnetic SI responses to electrical stimuli (Jousmäki & Forss, 1998; Torquati et al., 2002), including the P50m/P60m (Lin et al., 2003), which is seen as the direct magnetic analogue of the electrophysiological P50 (Wikström et al., 1996). Crucially however, our finding shows that the results achieved using electrical stimulation, beginning with Libet et al. (1967), may not hold generally, but may instead be peculiar to studies using this particular stimulation type. In other words, our finding suggests that somatosensory NCCs can vary depending on the type of stimulation used.

Microneurography studies (Johansson & Vallbo, 1979; Vallbo & Hagbarth, 1968; Vallbo & Johansson, 1984) showed that the sensory thresholds of type 1 rapidly adapting (RA) nerve fibers connected to Meissner corpuscles in the glabrous skin (Handler & Ginty, 2021; Vallbo & Johansson, 1984) and slowly adapting (SA) fibers in the hairy skin (Järvilehto et al., 1981) correspond to subjective detection thresholds, and that extremely sparse peripheral input to these receptors can be sufficient to induce conscious detection. Based on these results, Soininen and Järvilehto (1983) proposed that the tactile P50 response may be triggered by this peripheral activity volley and is related to conscious perception. However, some authors believe that potentials in this latency range are too early to directly reflect endogenous processing of the stimulus, and, to the extent that they are predictive of perceptual outcomes, rather reflect the prestimulus attentional state, anticipation of, and/or expectations about the stimulus (Desmedt & Tomberg, 1989; Josiassen et al., 1982). The P50/P60m has been proposed in part to reflect inhibitory processing (Wikström et al., 1996) that sharpens the initial thalamo-cortical input (itself reflected by the N20) with respect to (in-or extrinsic) noise (Nierhaus et al., 2015; Pleger & Villringer, 2013). While inhibitory processing is initiated already for subthreshold stimuli, it would then have to reach a sufficient level in order to enable conscious perception. This model matches well with studies reporting scaling of the P50 with stimulus intensity (Forschack et al., 2020), and it can also accommodate the enhancement of the P50 by attention (Desmedt et al., 1983; Forschack et al., 2017; Josiassen et al., 1982, 1990), which in this light appears as an attentional modulation of inhibitory networks in SI.

However, the question remains why the P50 amplitude is predictive of detection for tactile, but not electrical stimulation. An obvious difference between the two stimulation types lies in how they affect the physiological pathways leading from the index finger to SI: while the brief mechanical impulse of the piezoelectrically driven pin during tactile stimulation activates the Meissner’s corpuscles and, albeit to a lesser extent if at all, the Merkel cells in the fingertip, the effects of electrical stimulation are not limited to these receptor types, but will indiscriminately impact on all local fibers, bypassing receptors of all types. An electrically induced swath of activity arriving at SI may therefore be more likely than a mechanically induced swath to generate measurable inhibitory potentials already for subthreshold stimulation. Importantly, once the stimuli are perceived, there is visible scaling of P50 amplitudes with intensity for mechanical stimulation as well, both in our data and in two MEG studies that found an SI-generated M50 response to supra-threshold stimuli (Onishi et al., 2010; Yamashiro et al., 2019). There, the authors concluded that the M50 magnitude probably reflects the number of activated mechanoreceptors. Under the inhibitory model for the P50, this suggests the following picture: both with mechanical and electrical fingertip stimulation, the P50 scales with the number of activated local nerve fibers, and therefore with stimulus intensity, which directly influences the degree of inhibitory SI activity. In both cases, the subjective detection threshold coincides with the activation threshold of the type 1 RA fibers connected to Meissner type end organs (Johansson & Vallbo, 1979; Soininen & Järvilehto, 1983; Vallbo & Johansson, 1984). Therefore, in the case of mechanical stimulation, P50 scaling with stimulus intensity becomes apparent only once the intensity is high enough to cross this threshold, as only then is a P50 generated in the first place. In the case of electrical stimulation, on the other hand, a measurable P50 that scales with intensity emerges already at intensities below the type 1 RA fiber (and therefore, subjective detection) threshold, because some non-RA nerve fibers are already being activated. Essentially, in the case of electrical stimulation the binary detection effect of the P50 is masked by these subthreshold activations. When the median nerve at the wrist instead of the fingertip is stimulated, this masking phenomenon plays out even stronger, because an even greater variety of nerve types (including proprioceptive and afferent motor fibers) are targeted at this site (Koivikko, 1971). This may also help to explain why two studies found early activity to be predictive of detection despite using electrical stimulation (Hirvonen & Palva, 2016; Palva et al., 2005). Both these studies were conducted using MEG. Importantly, MEG is sensitive only to sources tangential to the scalp, that is, to activity coming predominantly from the cortical sulci, whereas EEG tracks activity from both gyri and sulci (M. Hämäläinen et al., 1993). If the above picture is correct, then EEG is likely to be more susceptible to the described masking effect, because it picks up more of the additional subthreshold SI activity induced by electrical stimulation than MEG does. Importantly, the suggested coincidence of the thresholds of subjective perception and type 1 RA unit activation whose crossing is partly reflected in the P50 does not necessarily mean that the P50 is an NCC proper, as opposed to a mere prerequisite of consciousness (Aru et al., 2012; de Graaf et al., 2012). It remains possible that more complex processing following the RA-mediated sharpening response in SI is necessary to render the stimulus conscious, as we discuss next.

### The N80-N140 complex

For mechanical but not electrical stimulation, we found a prominent N80 component for hits compared to misses over contralateral sensors C6 and FC6. A similar component has often been described in studies using strong electrical (Allison et al., 1992; Desmedt et al., 1983; W. R. Goff et al., 1962; Michie et al., 1987) or mechanical stimulation (Eimer & Forster, 2003; Guidotti et al., 2023; H. Hämäläinen et al., 1990; Schubert et al., 2008; Taylor-Clarke et al., 2002; Zopf et al., 2004). While the N80 was not present as a clearly separable component in the electrical group in our data, it is noteworthy that the contralateral part of the electrical N140 deflection began already around 60 ms, at the same time as the tactile N80, and was slightly bimodal, i.e., had a small peak already before the main peak at 133 ms. Interestingly, an occasionally bimodal N140 has been reported already by Goff et al. (1962), who compared SEPs to supra-threshold stimulation of the median nerve at the wrist and the index finger, as well as by Hämäläinen et al. (1990), who used mechanical stimulation of the middle finger. It is also discernible in somatosensory masking (Schubert et al., 2006) and peri-threshold detection studies (Forschack et al., 2020; Schröder et al., 2021), but has usually gone unnoticed. In some of these studies, the first of the two peaks is strongly reminiscent of the effect at 80 ms reported by Auksztulewicz and colleagues (2012; 2013). It is well known that the peak latency of the N140 is rather variable, and it has also been shown that it is a complex component containing several subcomponents (Allison et al., 1992; García-Larrea et al., 1995; Zopf et al., 2004), potentially reflecting different processes. It is therefore possible that the N140 generally contains an earlier N80-like part that is difficult to separate from the N140 proper when electrical stimulation is used, but becomes obvious with mechanical stimulation. Allison et al. (1992) observed that the electrically-induced N70 is only recordable in isolation at rather short ISIs, and “appears to be obscured by other long-latency activity” (p. 311) for longer ISIs, as they are typically used in NCC studies (including the present one). The MEG study by Jones et al. (2007), who used tactile threshold stimulation and investigated the response time course of a dipole localized to contralateral SI using a biophysical model, found that the earliest deflection predictive of subjective detection was a negative M70 component which in fact peaked at 80 ms (sometimes considerably later), and transitioned into a positive-going M135 deflection. Both responses were likely to be driven by excitatory feedback input from SII, with the M135 requiring additional thalamic input, which in turn was presumably driven by cortico-thalamic feedback, again likely involving SII. Based on these modelling results, Jones et al. (2007) believe that the M70 probably corresponds to the intracortically recorded N1 component found in monkeys for hits compared to misses in a detection task for both mechanical and electrical stimulation (Cauller & Kulics, 1991). There, the difference between the two stimulation types was that electrical stimulation activated a much larger horizontal (intra-layer) area of SI than mechanical stimulation, whereas the mechanically evoked N1 was more focal but also had a larger peak amplitude. These characteristics seem to match well with those of the scalp-level signals recorded in our study, where the tactile N80 peak is very prominent, while the corresponding first N140 peak during electrical stimulation is less discernible, and appears as part of the N140 component. Most likely then, our tactile N80 corresponds to the M70 of Jones et al. (2007) and the N1 of Cauller and Kulics (1991), while the M135, into which the M70 transitions, corresponds to the N140 proper. Given that the detection-predictive M70/N1/N80 seems to be heavily driven by cortico-cortical and cortico-thalamo-cortical feedback loops involving sensory cortices, and Auksztulewicz and colleagues (2012; 2013) have provided evidence for a complex origin of the N140 involving recurrent processing between SI and SII, the N80/N140 complex would be a natural candidate for an NCC according to recurrent processing theory (Lamme, 2006, 2010; Lamme & Roelfsema, 2000), which posits precisely this feature as a hallmark of consciousness.

*P100.* While the P100 and its magnetic analogues have been described in many studies, others reported not finding this component, and it has received various interpretations both in terms of its cortical origin and its functional significance. It can be elicited by mechanical (Eimer & Forster, 2003; H. Hämäläinen et al., 1990; Jones et al., 2007; Yamashiro et al., 2019), airpuff (Forss, Salmelin, et al., 1994) and electrical (Allison et al., 1992; G. D. Goff et al., 1977; Hari et al., 1983, 1984) stimulation of the median nerve both at the wrist and fingers (Desmedt et al., 1983; W. R. Goff et al., 1962). Its origins have been ascribed to area 1 of SI (Allison et al., 1992) and posterior parietal cortex (Desmedt & Tomberg, 1989; Forss, Hari, et al., 1994; Forss, Salmelin, et al., 1994), but most often to bilateral SII (Eimer & Forster, 2003; H. Hämäläinen et al., 1990; Hari et al., 1983, 1984; Uemura et al., 2021; Yamashiro et al., 2019). It can have large intra- and intersubject variability (W. R. Goff et al., 1962), and is sometimes found in only a fraction of participants (Desmedt & Robertson, 1977; Zhang & Ding, 2009). The P100 is clearly responsive to manipulations of selective attention (Desmedt et al., 1983; Desmedt & Tomberg, 1989; Eimer & Forster, 2003; Josiassen et al., 1982; Michie et al., 1987), but it has also been related to conscious perception (Jones et al., 2007; Schubert et al., 2006), which on one leading account is itself dependent on attention as a necessary prerequisite (Dehaene & Naccache, 2001). In our study, we found a main effect of detection for the P100, which appears to be more strongly driven by the tactile group, but is still visible in the electrical group (Fig. 4). This is in line with the masking study using electrical stimuli by Schubert et al. (2006) and the threshold detection study by Jones et al. (2007) using mechanical stimuli, but at odds with several other threshold detection studies that used electrical stimulation and reported little or no trace of such an effect on the P100 (Auksztulewicz et al., 2012; Auksztulewicz & Blankenburg, 2013; Schröder et al., 2021). This divergence may be explained by the different stimulation site: whereas these studies targeted the median nerve at the wrist, Jones et al. (2007) stimulated the middle finger, and our study and the one by Schubert et al. (2006) stimulated the index finger. Goff et al. (1962) note that median nerve stimulation evoked larger, more distinct potentials than finger stimulation; and while the P100 in Schröder et al. (2021) was mostly driven by stimulus intensity, detection probability explained part of this component as well, suggesting that processes related to somatosensory awareness contribute to the P100 with median nerve stimulation as well, even though this may be harder to detect in this case.

*P300*. Unlike the other components, the P300 is a supramodal, task-related component that presumably reflects multiple processes and has received many functional interpretations (Verleger, 2020). In the context of NCC research, it has been considered a hallmark of conscious processing in the framework of Global Neuronal Workspace theory (Dehaene & Changeux, 2011; Dehaene & Naccache, 2001) until more recent studies showed that it reflects post-perceptual processes more closely (Dellert et al., 2021; Pitts et al., 2014; Schlossmacher et al., 2021; Schröder et al., 2021). In the somatosensory modality, it is elicited by both mechanical and electrical stimulation (G. D. Goff et al., 1977; Soininen & Järvilehto, 1983) and has long been known to reflect the processing of rare target stimuli (Desmedt et al., 1977; Desmedt & Robertson, 1977; Josiassen et al., 1982, 1990; Michie et al., 1987; Nakajima & Imamura, 2000). In threshold detection tasks, it reflects somatosensory awareness both categorically (Auksztulewicz et al., 2012) and parametrically (Auksztulewicz & Blankenburg, 2013), but ceases to do so when post-perceptual processes are controlled with a matching task (Schröder et al., 2021). Our study is at odds with this latter finding, in that we find larger P300 amplitudes for hits compared to misses in both the tactile and electrical groups, leading to a significant main effect of detection for this component that, unlike the P50 effect, was driven equally strongly by both groups. Schröder et al. (2021) stimulated the left median nerve at the wrist instead of the tip of the left index finger, where 1) fewer nerves and nerve types are targeted and 2) threshold intensities are much lower (1.46 mA in our study versus 2.53 mA in the matching task and 2.88 mA in the direct report task in Schröder et al., 2021). Nakajima & Imamura (2000) have shown that the somatosensory-evoked P300 has (at least) an endogenous and an exogenous component, in that its amplitude is modulated by attention, but also stimulus intensity. Also note that the P300 amplitude in the matching task in Schröder et al. (2021) is clearly higher for hits than misses in the subsample, although this difference was not formally analyzed in order to avoid double dipping. Conversely, despite the main effect of detection in our study, the P300 difference between hits and misses is not nearly as pronounced as it appears to be in the direct report task in Schröder et al (2021). Our results thus suggest that conscious processing still contributes to the P300 to some degree even when post-perceptual processes are controlled for, but that other factors (in particular, stimulus intensity and the additional input in the case of wrist stimulation, e.g., from proprioceptive and afferent motor fibers) can override this contribution. While this qualifies the idea that the P300 is entirely unrelated to conscious perception, it does by no means challenge the important insight that post-perceptual processes are the main drivers of this component and need to be controlled for in NCC studies.

## Conclusion

In conclusion, our study demonstrates that the somatosensory NCCs strongly depend on the type of stimulation used. Most importantly, the P50 component, which has until now been considered to reflect physical stimulus attributes, is part of the NCCs (at the very least in the broad sense which encompasses prerequisites and consequences of consciousness as well) under physiological conditions, i.e., when using “naturalistic” tactile as opposed to “artificial” electrical stimulation. Further, our results corroborate the N140 as a robust NCC with both types of stimulation, with a pronounced N80-N140 partition in the tactile case that is less evident in the electrical case. Finally, the P300 effect of perception in our data suggests that the degree to which this component reflects perceptual or post-perceptual processes, respectively, may depend on the stimulation site (e.g., wrist vs. finger), once again illustrating the susceptibility of somatosensory NCCs to different stimulation protocols.

## Author contributions

Jona Förster (Conceptualization, Investigation, Formal analysis, Methodology, Software, Visualization, Writing – original draft, Writing – review and editing), Giovanni Vardiero (Investigation, Writing – review and editing), Till Nierhaus (Conceptualization, Supervision, Writing – review and editing), Felix Blankenburg (Conceptualization, Supervision, Methodology, Writing – review and editing).

## Data availability

Preprocessed data and analysis scripts are available from the corresponding author upon request.

